# LRSDAY: Long-read Sequencing Data Analysis for Yeasts

**DOI:** 10.1101/184572

**Authors:** Jia-Xing Yue, Gianni Liti

## Abstract

Long-read sequencing technologies have become increasingly popular in genome projects due to their strengths in resolving complex genomic regions. As a leading model organism with small genome size and great biotechnological importance, the budding yeast, *Saccharomyces cerevisiae*, has many isolates currently being sequenced with long reads. However, analyzing long-read sequencing data to produce high-quality genome assembly and annotation remains challenging. Here we present LRSDAY, the first one-stop solution to streamline this process. LRSDAY can produce chromosome-level end-to-end genome assembly and comprehensive annotations for various genomic features (including centromeres, protein-coding genes, tRNAs, transposable elements and telomere-associated elements) that are ready for downstream analysis. Although tailored for *S. cerevisiae*, we designed LRSDAY to be highly modular and customizable, making it adaptable for virtually any eukaryotic organisms. Applying LRSDAY to a *S. cerevisiae* strain takes ∼43 hrs to generate a complete and well-annotated genome from ∼100X Pacific Biosciences (PacBio) reads using four threads.

## INTRODUCTION

Twenty years ago, the genome sequence of the budding yeast *Saccharomyces cerevisiae* was published. As the first complete eukaryotic genome ever sequenced, this marked a major scientific milestone in biology. Since then, the genomes of many model and non-model organisms have been sequenced especially after the emergence of next-generation sequencing (NGS) technologies. Despite the notably improved throughputs, these NGS technologies suffer from the limitation of short reads and usually result in highly fragmented genome assemblies containing numerous gaps and local mis-assemblies. The recently developed long-read sequencing technologies represented by Pacific Biosciences (PacBio) and Oxford Nanopore offer compelling alternatives to overcome such hurdles, producing high-quality genome assemblies with significantly improved continuity and accuracy. Although initially tested in microbial genome sequencing, their recent applications in complex mammalian and plant genomes also achieved very impressive results^1–4^. With such new sequencing technologies, challenging genomic regions with enriched repetitive elements, strongly biased GC%, or complex structural variants can often be correctly resolved. It is therefore anticipated that genome sequencing projects will routinely adopt long-read-based sequencing technologies in the coming years.

Yeast is a leading model organism with great importance in both basic biomedical research and biotechnological applications. Its small genome size makes it particularly suitable for long-read-based high-coverage genome sequencing. The resulting complete genome assembly with fully-resolved subtelomere structure can in turn illuminate the genetic basis of many complex phenotypic traits with unprecedented resolution. Recently, we used the long-read sequencing technologies to generate the first panel of population-level end-to-end reference genomes of 12 yeast strains representing the major subpopulations of the partially domesticated *S. cerevisiae* and its sister species *Saccharomyces paradoxus*^5^. In addition, there have been a number of other studies carrying out long-read sequencing for many *S. cerevisiae* strains^6–9^. Given the vast genomic and phenotypic diversity of *S. cerevisiae*, we expect the incoming collection of long-read-based high-quality genome assemblies of strains from widespread geographic locations and ecological niches will substantially deepen our understanding on the *S. cerevisiae* natural genetic variation and its associated biotechnological values.

Here we present a highly organized and modular computational framework named Long-Read Sequencing Data Analysis for Yeasts (LRSDAY), which enables effortless high-quality yeast genome assembly and annotation production from raw long-read sequencing data. Although a few popular long-read-based genome assemblers^10–12^ and one yeast-specific gene annotation tools^13^ have been developed, a single integrated solution that handles the entire process in a seemless way is still much needed. Toward this goal, we revamped our original workflow for deriving the yeast population-level reference genomes^5^ into a highly streamlined and self-contained package with modular design and automated implementation. Under the hood, LRSDAY contains a series of task-specific modules handling long-read-based *de novo* genome assembly, short-read-based assembly polishing, reference-guided assembly scaffolding, as well as comprehensive genomic feature annotations. These tasks can be run individually, selectively or coordinately depending on users’ needs. LRSDAY supports both leading long-read sequencing technologies: PacBio and Oxford Nanopore. Running the full LRSDAY workflow, the final output is a chromosome-level genome assembly with high-quality annotations of centromeres, protein-coding genes, tRNAs, transposable elements (TEs; Ty1-Ty5 for yeasts), and telomere associated core X and Y’ elements. LRSDAY is shipped with various auxiliary scripts, configuration files and supporting data that enable its semi-automatic installation, configuration, and execution with minimal manual intervention. This design concept greatly alleviates the technical barrier for bench biologists with limited bioinformatics experiences (Box 1). In addition, a real case example and its final outputs are also provided for users’ test and comparison. All these task-specific modules, auxiliary files, installed tools, sample outputs, together with the user-created project directories for the testing example and users’ own data are hosted under the same home directory ($LRSDAY_HOME) in a self-contained fashion (Fig. 1). This design makes LRSDAY well-isolated from the rest of the system and therefore greatly improved its portability. To sum up, LRSDAY is a highly transparent, automated and powerful computational framework that handles both genome assembly and annotation, which suits the needs of the yeast community in the era of long-read sequencing.

**Figure 1.**
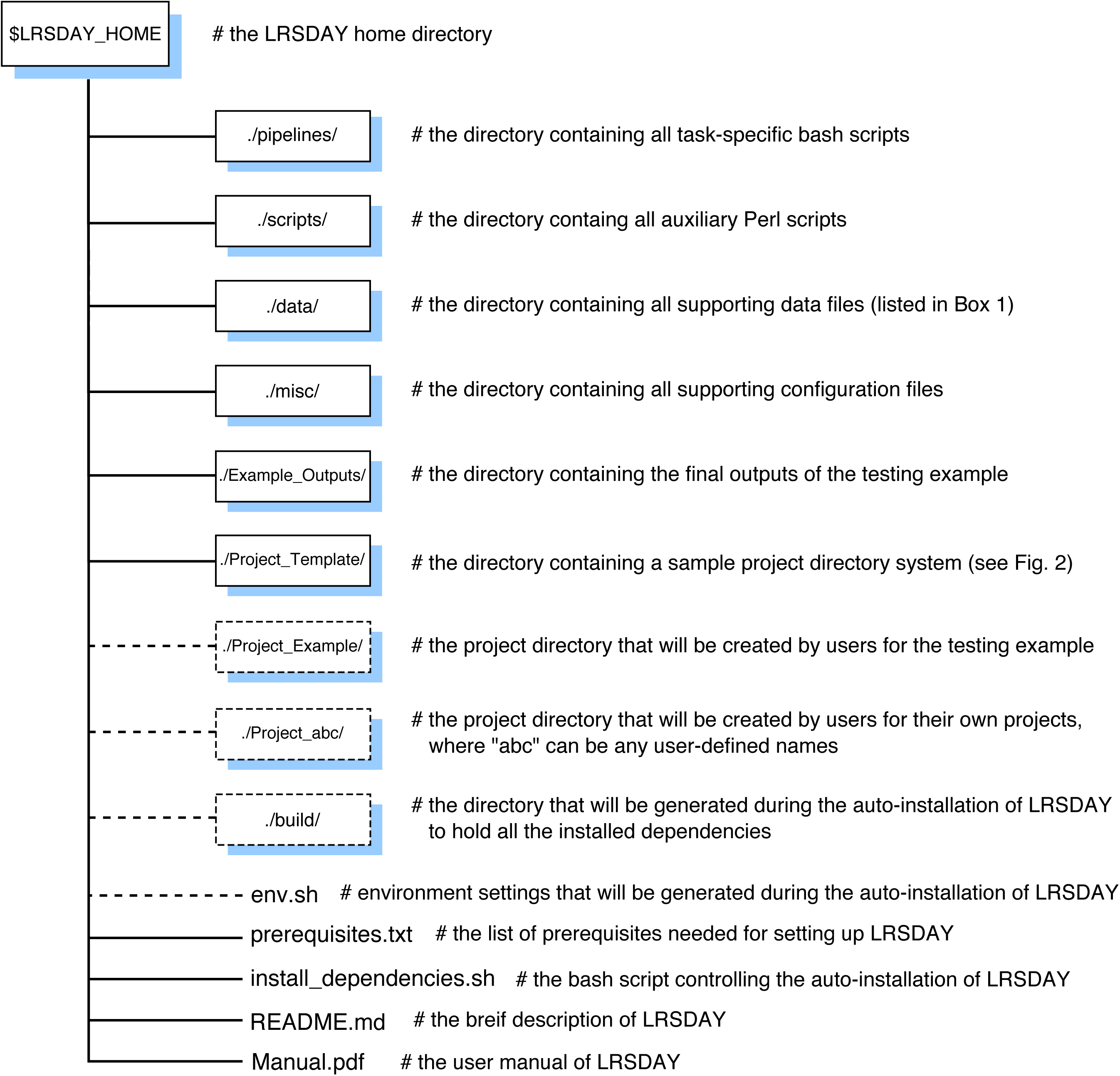
Overview of the LRSDAY directory system. All the top-level directories (boxes, solid lines) and individual files of LRSDAY are listed and briefly described. Additional directories and files will be generated during the installation or execution of LRSDAY (boxes, dashed lines).

### Overview of the LRSDAY workflow

Genome assembly and annotation are complex computational processes with many intermediate steps and inputs/outputs involved. With LRSDAY, we designed a highly structured project directory system to help users to run the whole workflow in an organized and modular way (Fig. 2). Within such project directory system, the three subdirectories holding the pre-shipped reference genome as well as the user-supplied long (PacBio or Oxford Nanopore) and short (Illumina) reads are numbered as “00” and the task-specific subdirectories are numbered sequentially from “01” to “13” according to their execution orders. For each subdirectory, a self-explained name is attached after the number index to help users to navigate through the whole workflow. To run each task, users only need to edit (e.g. to specify the input and output file paths) and execute the task-specific pipeline scripts pre-placed in these subdirectories. These pipeline scripts will automatically set environment variables, process the data, and formulate the results. All computationally intensive tasks can be processed using multiple threads to substantially save computation time. Below we briefly describe the computational processes underneath each task-specific module:

**Figure 2.**
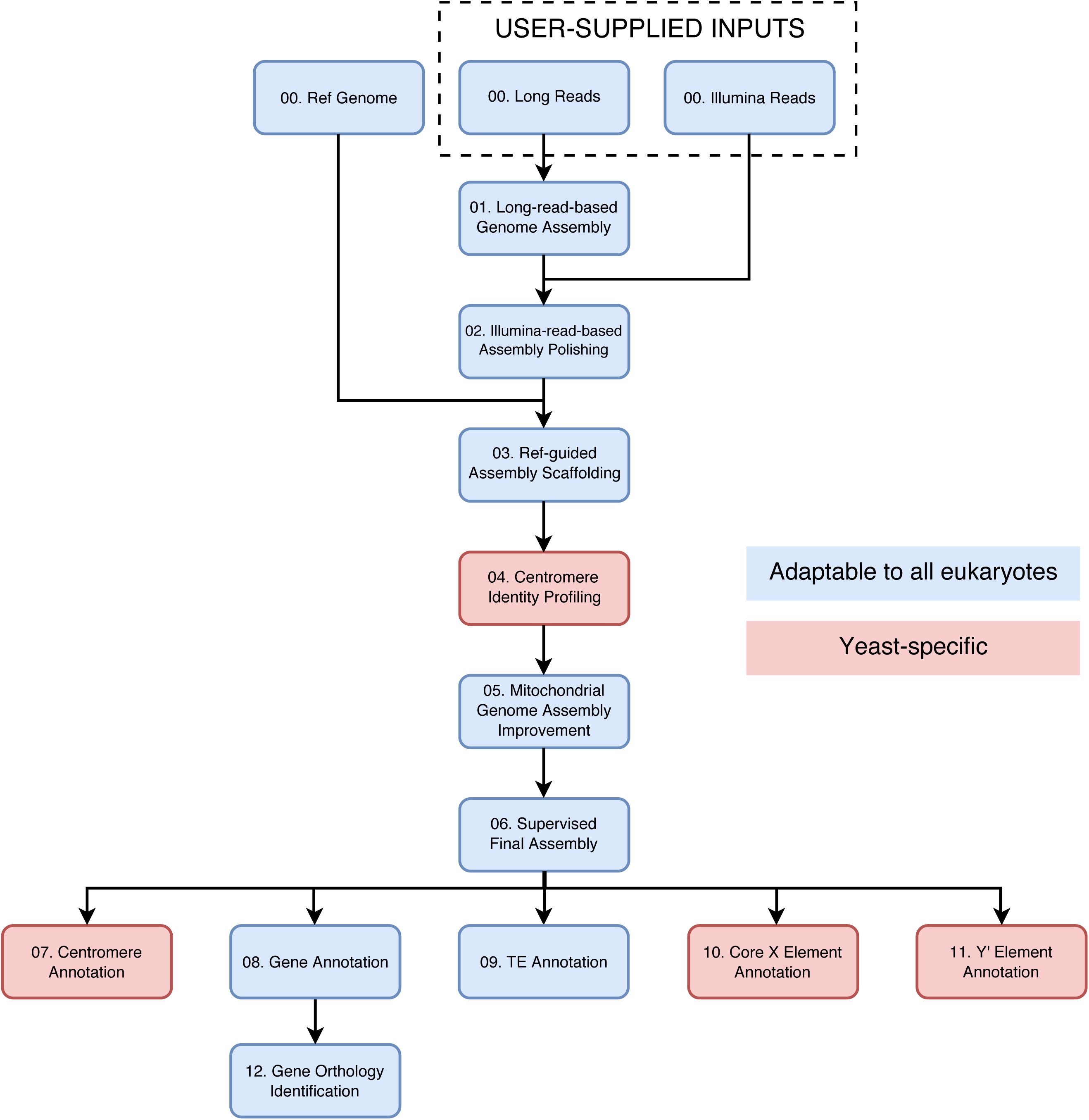
The workflow of LRSDAY. Each box represents an individual module. These modules are numbered according to their processing order in LRSDAY. Modules that can be adapted for other eukaryotes are colored in light blue while those yeast-specific ones are colored in light red.

**01. Long-read-based_Genome_Assembly**: Long reads generated from PacBio or Oxford Nanopore technologies are used to perform *de novo* genome assembly using an overlap-layout-consensus (OLC) algorithm.

**02. Illumina-read-based_Assembly_Polishing**: Illumina reads are first clipped and trimmed to remove potential sequencing adapters and low-quality regions. The cleaned reads are subsequently mapped to the raw long-read-based genome assembly. The resulting bam file is further processed for alignment sorting, mate information and read group fixing, duplicates removal as well as local realignment. Finally, the processed bam file is used for correcting base-level errors of the long-read-based assembly.

**03. Reference-guided_Assembly_Scaffolding**: The contigs from the polished genome assembly are first aligned to the reference genome to identify their shared sequence homology, based on which reference-guided assembly scaffolding is subsequently performed. The chromosomal identity of each scaffold is labeled accordingly. Structural rearrangements captured in the original contigs will stay untouched during the scaffolding.

**04. Centromere_Identity_Profiling**: The pre-shipped *S. cerevisiae* centromere sequences are searched against the scaffolded assembly for chromosome-specific centromere identity profiling.

**05. Mitochondrial_Genome_Assembly_Improvement**: The polished contigs corresponding to the mitochondrial genome are re-collected from the scaffolded assembly. The mitochondrial contigs spanning over the designated starting point (the *ATP6* gene by default) are broken into subsegments to prevent assembly problems caused by the circular organization of the mitochondrial genome. The resulting contigs are then re-assembled into a single linear sequence, which is further circularized by the designated *ATP6* starting point. The nuclear scaffolds and the circularized mitochondrial sequence together form the improved genome assembly.

**06. Supervised_Final_Assembly**: A modification list containing the ordering, orientation and naming information of each sequence from the improved genome assembly is generated for users to review and to make manual adjustment when needed. The final genome assembly is further generated based on the user-edited modification list.

**07. Centromere_Annotation**: The pre-shipped *S. cerevisiae* centromere sequences are searched against the final genome assembly for centromere annotation.

**08. Gene_Annotation**: *De novo* protein-coding and tRNA gene annotations are performed for the final genome assembly, which are further leveraged by the expressed sequence tags (ESTs) and protein sequences alignments.

**09. TE_Annotation**: The pre-shipped curated TE library (containing the LTRs and internal sequences of the *S. cerevisiae* and *S. paradoxus* Ty1-Ty5 by default) is searched against the final genome assembly to identify TEs. The identified TEs are further curated and classified into the full-length, truncated, and solo-LTRs of Ty1-Ty5.

**10. Core_X_Element_Annotation**: The pre-shipped curated hidden Markov model (HMM) of the *S. cerevisiae* core X elements is searched against the final genome assembly to annotate core X elements.

**11. Y_Prime_Element_Annotation**: The pre-shipped representative *S. cerevisiae* Y’ element sequence is searched against the final genome assembly to annotate Y’ elements. Note that Y’ element can have long, short or degenerated forms^14^, and we used a representative long-form Y’ element as the query to maximize the power of our Y’ element annotation.

**12. Gene_Orthology_Identification**: The annotated protein-coding genes are compared with the reference protein-coding genes based on both sequence homology and gene order conservation to identify gene orthology relationship between these two sets. Based on such gene orthology relationship, the *Saccharomyces* Genome Database (SGD;http://www.yeastgenome.org/) systematic names are assigned to the annotated protein-coding genes.

**13. Annotation_Integration**: The annotations of centromeres, TEs, protein-coding genes, tRNAs, as well as core X and Y’ elements are combined and sorted to form a final integrated multi-feature annotation.

### Limitations and potential adaptation

In its distributed form, LRSDAY is tailored for the model budding yeast *S. cerevisiae* and its closely related sister species *S. paradoxus* with a number of pre-shipped auxiliary data files configured accordingly. However, given its modular design, the backbone of LRSDAY can be adapted for virtually any eukaryotic organisms to perform genome assembly, assembly polishing, reference-guided scaffolding, protein-coding genes and tRNA annotations, gene orthology identification, and annotation integration. Moreover, those assembly polishing, scaffolding and various annotation modules (Task 02-13) can also be used to analyze existing genome assemblies derived from any or any combination of sequencing technologies. Such flexibility makes LRSDAY very useful for expanded use cases and therefore suits the needs of a broader audience.

### Expected improvements

As thousands of yeast strains have been or are currently under sequencing^15–17^, our knowledge in the overall genome content diversity^18^ of this important model organism is expanding rapidly, revealing a whole new picture of the pan-genome diversity of *S. cerevisiae*. For example, our lab is currently working on characterizing the pan-genome of >1,000 *S. cerevisiae* isolates across the globe (http://1002genomes.u-strasbg.fr/). Future developments of LRSDAY will incorporate such pan-genome dataset to provide additional annotation information for those non-reference genes, especially with regard to their evolutionary origin, population prevalence, and putative functions. Such information will greatly help users to dissect and interpret complex genotype-phenotype interactions in diverse ecological and biotechnological settings. In the protocol described below, we provide a step-by-step walkthrough on how to install, configure, and run LRSDAY for our prepared testing example. Further tips on adapting LRSDAY to other eukaryotic organisms can be found in Box 2.

## MATERIALS

### EQUIPMENT

#### Hardware, operating system and network

- This protocol is designed for a desktop or computing server running with x86-64-bit Linux operating system. Multithreaded processors are preferred to speed up the process since many steps can be configured to use multiple threads in parallel. For assembling and analyzing the budding yeast genomes (genome size = ∼12 Mb), at least 16 Gb of RAM and 100 Gb of free disk space are reccomended. When adapted for other eukaryotic organisms with larger genome sizes, the RAM and disk space consumption will scale up, majorly during *de novo* genome assembly (performed by Canu^12^). Plese refer to Canu’s manual (http://canu.readthedocs.io/en/latest/tutorial.html) for suggested RAM and disk space consumption for assembling large genomes. Stable Internet connection is required for the installation and configuration of LRSDAY as well as for retrieving the test data.

#### Software or library requirements

- Bash (https://www.gnu.org/software/bash/)
- Bzip2 (http://www.bzip.org/)
- Cmake (https://cmake.org/)
- GCC and G++ v4.7 or newer (https://gcc.gnu.org/)
- Git (https://git-scm.com/)
- GNU make (https://www.gnu.org/software/make/)
- Gzip (https://www.gnu.org/software/gzip/)
- Java runtime environment (JRE) v1.8.0 or newer (https://www.java.com)
- Perl v5.12 or newer (https://www.perl.org/)
- Python v2.7.9 or newer (https://www.python.org/)
- Python v3.6 or newer (https://www.python.org/)
- Tar (https://www.gnu.org/software/tar/)
- Virtualenv v15.1.0 or newer (https://virtualenv.pypa.io)
- Wget (https://www.gnu.org/software/wget/)

#### Input data

- Long reads: A single FASTQ file containing PacBio or Oxford Nanopore reads is needed, which will be used for long-read based *de novo* genome assembly (Task 01).
- Short reads: If paired-end Illumina sequencing is performed, two FASTQ files containing the forward and reverse Illumina reads respectively are needed. If only single-end Illumina sequencing data is available, one FASTQ file containing the single-end reads is needed. The short-read sequencing reads will be used for assembly polishing (Task 02).
- Reference genome: For the budding yeast *S. cerevisiae*, we pre-shipped two reference genome files (one original assembly and one with subtelomeres and chromosome-ends hard-masked based on our previous study^5^). The masked version is used for chromosomal scaffolding to minimize the confounding effect due to interchromosomal subtelomeric rearrangemnets. When working with organisms of which the subtelomeric regions are undefined, users can just use a single raw reference genome instead. The reference genome file(s) will be used for reference-guided scaffolding, mitochondrial genome assembly improvement and supervised final genome assembly (Task 03, 05 and 06 respectively).
- A number of *S. cerevisiae*-specific auxiliary data have been pre-shipped with LRSDAY for genomic feature annotation and gene orthology identification (Task 07-12).

### Example data

- The test run sequencing data comes from our recent study^5^, which consists of both PacBio and Illumina reads produced from the *S. cerevisiae* strain SK1. The PacBio reads can be retrieved with the ENA analysis accession number ERZ448251. The Illumina reads can be retrieved with the SRA sequencing run accession number SRR4074258. The *S. cerevisiae* reference genome files used by LRSDAY is taken from the same study with the Genbank accession number GCA_002057635.1. For the testing example, we have provided bash scripts to automatically download and setup these data in LRSDAY.

## PROCEDURE

### *Download, install and configure LRSDAY*• **TIMING <1 h**

1) Enter the following commands in a terminal window:

$ git clone https://github.com/yjx1217/LRSDAY.git

$ cd LRSDAY

$ bash install_dependencies.sh

**▴CRITICAL STEP** Make sure you have a fast and stable internet connection when running this step since many tools will be downloaded here. Check that all the prerequisites (see this protocol or the “prerequisite.txt” file in the downloaded LRSDAY directory) have been installed on your system.

**▴CRITICAL STEP** Upon the successful completion of executing the bash script, you should see a confirmation message prompted out: “LRSDAY message: This bash script has been successfully processed! :)”. Otherwise, it means an error has occurred during the execution of the bash script, which interrupted the automatic installation process. This also apply to all the bash scripts used in Step 25-45. Whenever an error is encountered, please check the error message and refer to the troubleshooting section when available. When the cause of the error is fixed, it is always recommended to completely delete the output files generated by the old run before re-initiate the new run.

**? TROUBLESHOOTING**

2) Load the environment settings for LRSDAY by entering:

$ source env.sh

After loading the pre-configured environment settings, the current directory should be assigned to the environment variable $LRSDAY_HOME. You can check to see if the full path to your current directory is displayed after entering:

$ echo $LRSDAY_HOME

**▴CRITICAL STEP** Make sure to run this command to load the pre-configured environment settings before the manual setup of LRSDAY in Step 3-22. If you exited your terminal session before or in the middle of your manual setup, you need to re-load the environment settings before proceeding. These environment settings will be automatically loaded each time the task-specific bash pipelines of LRSDAY are executed.

**? TROUBLESHOOTING**

3) Pay attention to the final message produced by the install_dependencies.sh script (Step 1) on manual setup instructions. Although most required tools have been automatically installed and configured, manual downloading and/or configuration are needed for GATK^19^, RepeatMasker^20^ and MAKER^21^ due to license restriction of these tools or of their dependent databases. Below, we will go over this process in Step 4-22.

4) Go to the official website of GATK (https://software.broadinstitute.org/gatk/download) and download GATK-3.8 after logging in. Registration is needed for unregistered GATK users.

5) Place the downloaded GATK package (file name: “GenomeAnalysisTK-3.8-0.tar.bz2”) in the “LRSDAY/build/GATK-3.8” directory by entering the following command in terminal:$ mv GenomeAnalysisTK-3.8-0.tar.bz2 $LRSDAY_HOME/build/GATK-3.8

6) Uncompress and set up GATK by entering:$ tar xjf GenomeAnalysisTK-3.8-0.tar.bz2

$ mv GenomeAnalysisTK-3.8-0*/GenomeAnalysisTK.jar.$ chmod 755 GenomeAnalysisTK.jar

7) Go to the official website of MAKER (http://yandell.topaz.genetics.utah.edu/cgi-bin/maker_license.cgi) and download MAKER v3.00.0-beta” after registration.

8) Place the downloaded file (named as “maker-3.00.0-beta.tgz”) in the “LRSDAY/build/” directory by entering the following command in terminal:$ mv maker-3.00.0-beta.tgz $LRSDAY_HOME/build/

9) Uncompress MAKER by entering:$ tar xzf maker-3.00.0-beta.tgz

10) Relocate to the “./maker/src” subdirectory by entering:$ cd./maker/src

11) Replace the “Build.PL” script with the LRSDAY pre-shipped copy:$ cp $LRSDAY_HOME/misc/maker_Build.PL Build.PL

12) Run the replaced “Build.PL” script to configure MAKER:$ perl Build.PL

13) The Build.PL script will prompt prerequisite warnings/errors and a question related to “MPI”. Here, you should not see any required Perl library that is not installed. You might see the recommended Perl library “DBD::Pg” is not installed, which is fine. You can safely ignore the warnings for all required external programs. Accept the default answer to the MPI question by just pressing enter.

**? TROUBLESHOOTING**

14) Finalize the installation by entering:$./Build install

15) Relocate to the “$LRSDAY_HOME/build/RepeatMasker” directory :$ cd $LRSDAY_HOME/build/RepeatMasker

16) Go to the website (http://tandem.bu.edu/trf/trf409.linux64.download.html) to download trf v4.09. Place the downloaded file (“trf409.linux64”) in the “$LRSDAY_HOME/build/RepeatMasker” directory. Then, change the access permissions and create a symbolic link for this file by entering:

$ chmod 755 trf409.linux64

$ ln -s trf409.linux64 trf

17) Go to the website (http://www.girinst.org) and register a user account to download the most recent version of the RepeatMasker-edition RepBase database (current version: RepBaseRepeatMaskerEdition-20170127) for RepeatMasker. Place the downloaded file in the “$LRSDAY_HOME/build/RepeatMasker” directory.

18) Uncompress the RepBase database:

$ tar xzf RepBaseRepeatMaskerEdition-*.tar.gz

19) Get the full path to the current directory by entering:

$ pwd

Remember this path since it will be used for Step 22.

20) Get the full path to $blast_dir by entering:$ echo $blast_dir Remember this path since it will be used for Step 22.

**? TROUBLESHOOTING**

21) Run the configure script for RepeatMasker:

$ perl./configure

22) The configure script above will prompt several questions:

- Enter “env” for the question about the installation path of Perl.
- Just press enter for the question about the installation path of RepeatMasker.
- Enter the path that you obtained in Step 19.
- Enter “2” for the question about selecting a search engine. Then enter the path that you obtained in Step 20.
- Just press enter for the question about the default search engine.
- Enter “5” to finalize the configuration.

### *Run LRSDAY with a testing example* **• TIMING _42 h_**

Before proceeding to your own project, it is advised to first run our prepared testing example to check if LRSDAY is working properly as well as to get acquainted with the logic and workflows of LRSDAY.

23) Go to the $LRSDAY_HOME directory by entering:

$ cd $LRSDAY_HOME

24) Create the project directory. When running LRSDAY with your own data, it is recommended to make a copy of our “Project_Template” directory to create your own project directory such as “Project_abc”, where “abc” can be any string containing letters, numbers, or underscores. For this test example, we will make a copy of the “Project_Template” directory and named it as “Project_Example” by entering:

$ cp -r Project_Template Project_Example

25) Prepare the reference genome files. When running LRSDAY with your own data, you can directly put the reference genome (in FASTA format) in the “00.Ref_Genomes” subdirectory of your project directory (e.g. “Project_abc”). For this testing example or if your sequenced organism is *S. cereviaie*/*S. paradoxus*, you can prepare the reference genome by entering:

$ cd./Project_Example/00.Ref_Genome

$ bash LRSDAY.00.Prepare_Sc_Ref_Genome.sh

26) Prepare the long reads. When running LRSDAY with your own data, you can directly put the long reads in the “00.Long_Reads” subdirectory of your project directory (e.g. “Project_abc”). For this testing example, you can download the long reads by entering:

$ cd./../00.Long_Reads

$ bash LRSDAY.00.Retrieve_Reads_from_ENA_BAM.sh

27) Prepare the Illumina reads. Like for the long reads, you can directly put your Illumina reads in the “00.Illumina_Reads” subdirectory of your project directory (e.g. “Project_abc”) when running LRSDAY for your own data. For this testing example, you can download the Illumina reads by entering:

$ cd./../00.Illumina_Reads

$ bash LRSDAY.00.Retrieve_Reads_from_SRA.sh

**? TROUBLESHOOTING**

28) Run long-read-based *de novo* genome assembly. This step can be run with multiple threads to speed up the process. Depending on the CPU configuration of your Linux server/desktop, you might want to edit the “threads=” option in the bash script “LRSDAY.01.Long-read-based_Genome_Assembly.sh” to enable multiple threading. You can do the same for all the following tasks whenever the “threads=” option is provided in the corresponding task-specific bash script. Simple text editors such as emacs, vim, gedit or pico are highly recommended for such editing. Rich text editors might not work. Upon the completion of genome assembly, a summary file (“SK1.canu.stats.txt” for this test example) will be generated to report some basic summary statistics (e.g. total assembly size, N50, L50, GC%, etc) to help you to gauge the genome assembly quality (Table 1).

$ cd./../01. Long-read-based_Genome_Assembly

$ bash LRSDAY.01.Long-read-based_Genome_Assembly.sh

**TABLE 1.** | Assembly statistics for *S. cerevisiae* SK1 assembled in the testing example.

| Assembly statistics | Raw assembly | Scaffolded assembly | Final assembly |
| --- | --- | --- | --- |
| Total sequence count | 22 | 20 | 20 |
| Total sequence length (bp) | 12282921 | 12292985 | 12261860 |
| Min sequence length (bp) | 24410 | 24410 | 24410 |
| Max sequence length (bp) | 1480192 | 1480198 | 1480198 |
| Mean sequence length (bp) | 558314.59 | 614649.25 | 613093.00 |
| Median sequence length (bp) | 558503.50 | 637681.50 | 637681.50 |
| N50 (bp) | 829956 | 923665 | 923665 |
| L50 | 6 | 6 |  |
| N90 (bp) | 462498 | 462493 | 462493 |
| L90 | 14 | 13 | 13 |
| GC% | 38.21 | 38.17 | 38.24 |
| N% | 0.00 | 0.08 | 0.04 |

**▴CRITICAL STEP** Note this step will take long to finish (see **TIMING**), so we recommend to run this step and all the other time-consuming steps using “nohup” (https://en.wikipedia.org/wiki/Nohup), which allows the process to continue running after you exit the terminal or logout from the server. As an example, you can run the bash script using nohup as follows:

$ nohup bash LRSDAY.01.Long-read-based_Genome_Assembly.sh >run_log.txt 2>&1&

**▴CRITICAL STEP** When running LRSDAY with your own data, modify the bash script to specify the file paths and the prefix for your input and output data (same for all following steps).

**? TROUBLESHOOTING**

29) Polishing genome assembly with Illumina reads. This step can be run with multiple threads.

$ cd./../02.Illumina-read-based_Assembly_Polishing

$ bash LRSDAY.02.Illumina-read-based_Assembly_Polishing.sh

**▴CRITICAL STEP** When running LRSDAY for your own data, if you performed PacBio sequencing and also have access to the PacBio SMRT Analysis software package (http://www.pacb.com/products-and-services/analytical-software/smrt-analysis/), we recommend to run the first-pass of assembly polishing based on the raw PacBio reads by using PacBio’s own Quiver/Arrow pipeline^10^ before proceeding to this step. Likewise, if you performed Oxford Nanopore sequencing, we recommend using nanopolish (https://github.com/jts/nanopolish) or equivalent tools to polish the genome assembly using the raw Nanopore reads first.

30) Chromosome-level scaffolding for the raw *de novo* assembly. This step can be run with multiple threads. Upon completion, a list of summary statistics (“SK1.ragout.stats.txt” for this test example) will be generated for the scaffolded assembly (Table 1).

$ cd./../03.Reference-guided_Assembly_Scaffolding

$ bash LRSDAY.03.Reference-guided_Assembly_Scaffolding.sh

**▴CRITICAL STEP** lease heck the enerated enome-wide otplot “SK1.ragout.filter.pdf” (Fig. 3a) to verify the correctness of chromosome assignment and applying manual adjustment if necessary in Step 32. When running LRSDAY with your own data, you might see one scaffold correspond to more than one reference chromosomes, which could be due to shared sequence homology between duplicated regions or interchromosomal rearrangements Both types of events can be correclty interpreted based on the genome-wide dotplot generated in this Step. In either case, LRSDAY can correctly assign chromosomal identity of the corresponding scaffold based on its centromere annotation in Step 31. Also please check the “SK1.ragout.agp” file for the details of reference-based scaffolding. When running LRSDAY for your own data, if you have strong evidence for mis-scaffolding based on prior knowledge or other experimental data (e.g. mate-pair libraries or chromosomal contact data), you can break up the corresponding ragout scaffolds back to contigs and re-joined them with corrected order using the pre-shipped Perl scripts “break_scaffolds_by_N.pl”, “join_contigs_by_N.pl” and “extract_region_from_genome.pl” in the “$LRSDAY_HOME/scripts” directory. A scenario for such use case is when the breakpoints of structural rearrangements are also the breakpoints of the genome assembly. In this case, the reference-based scaffolding will arrange the contigs according to the reference genome configuration and therefore un-do the genome rearrangement.

**Figure 3.**
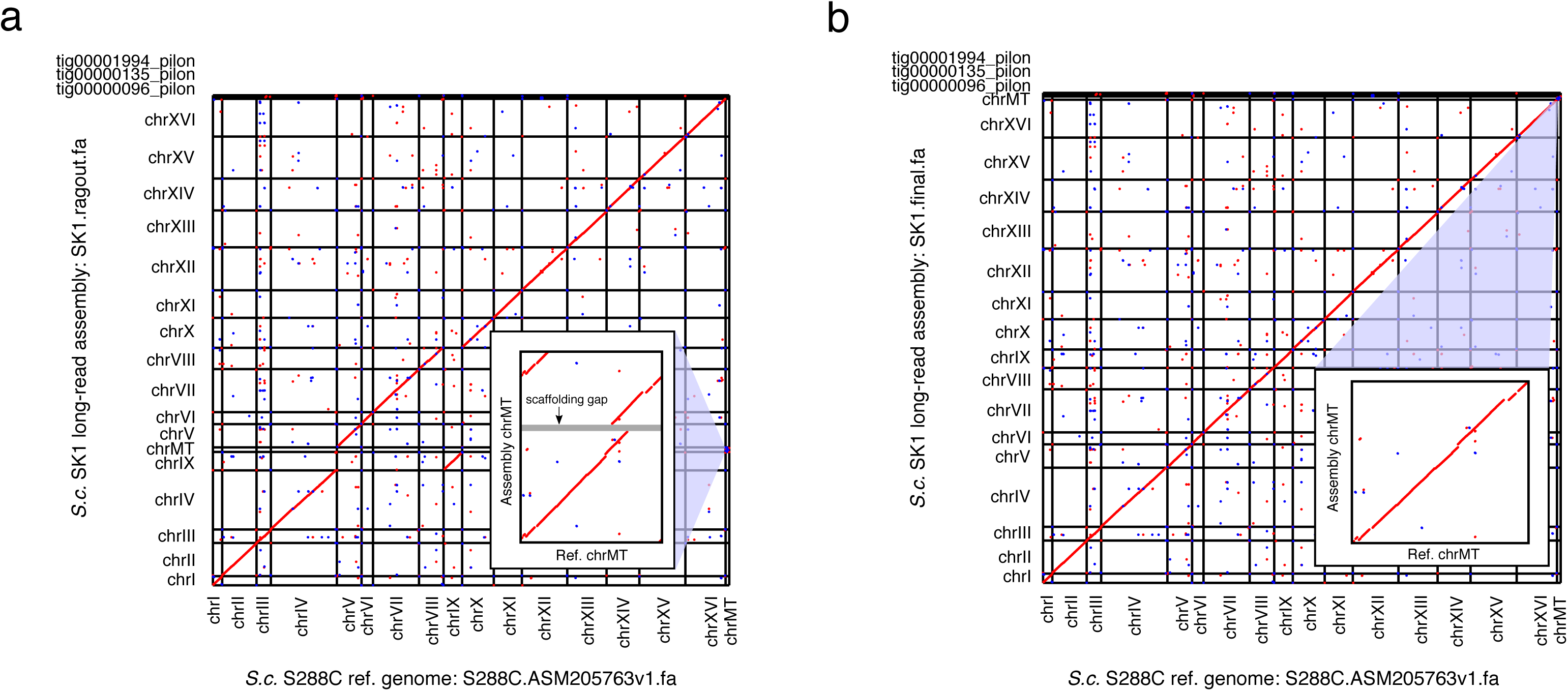
Genome-wide dotplots of the *S. cerevisiae* SK1 PacBio assembly generated in the LRSDAY testing example. Both the raw scaffolded assembly (panel a; generated at Step 30) and the final assembly (panel b, generated at Step 33) are analyzed. The forward and reverse sequence matches are depicted in red and blue respectively, while the zoomed-in views of the mitochondrial genome (chrMT) comparison are shown in insets. The scaffolding gap of the mitochondrial genome is indicated by the black arrow.

**▴CRITICAL STEP** Due to the high AT and repeat contents and the circular conformation of the mitochondrial genome, multiple contigs corresponding to the mitochondrial genome are often observed, as shown in the mitochondrial genome dotplot “SK1.ragout.chrMT.filter.pdf” (Fig. 3a, inset). A list of such mitochondrial contigs will be generated in the file “SK1.mt_contig.list”, which will be used in Step 32 for improving mitochondrial genome assembly.

31) Perform centromere profiling for the scaffolded genome assembly.

$ cd./../04.Centromere_Identity_Profiling

$ bash LRSDAY.04.Centromere_Identity_Profiling.sh

**▴CRITICAL STEP** The chromosome-specific centromere identies profiled here will be used as another layer of information for the final chromosomal identity assignment in Step 33. The profiled centromere idenitites should usually agree well with the chromosomal identity labeled in Step 30, so that chrI will have the CEN1 centromere and chrII will have the CEN2 centromere, etc. Exception can occur when interchromosomal rearrangements are involved in your sequenced species/strain. In such case, we recommend to name those rearranged chromosomes based on their encompassed centromeres in Step 33.

32) Mitochondrial genome assembly improvement.

$ cd./../05.Mitochondrial_Genome_Assembly_Improvement

$ bash LRSDAY.05.Mitochondrial_Genome_Assembly_Improvement.sh

**▴CRITICAL STEP** Check the “SK1.mt_improved.chrMT.filter.pdf” file for the final mitochondrial genome dotplot and compared with the mitochondrial genome dotplot generated in Step 30 to see if the mitochondrial genome assembly has been improved (Fig. 3b, inset).

33) Finalize the chromosome assignment and the genome assembly. This step consists of two substeps with manual intervention in between:

i. Generate the SK1.modification.list file. $ cd./../06.Supervised_Final_Assembly $ bash LRSDAY.06.Supervised_Final_Assembly.1.sh
ii. Edit the generated “SK1.modification.list” file based on the genome-wide dotplot generated in Step 30 and the centromere profiles generated in Step 31. The “SK1.modification.list” file consists of three comma-separated columns, which correspond to the original sequence name, sequence orientation, and new sequence name respectively. With this file, you can do three types of editing:
  a. If you need to change the current sequence order, you can move the corresponding row of that sequence upward or downward to reflect the correct order.
  b. If you need to invert the orientation of a given sequence, you can change its orientation from “+” to “-” in column 2.
  c. If you need to rename a given sequence, you can modify column 3 to specify the new name.

For the testing example here, we need to move the row “chrIX,+,chrIX” downward to place it after the row “chrVIII,+,chrVIII”, so that chrIX will be placed after chrVIII in the final assembly. Also, we need to change the row “chrMT_Contig1,+,chrMT_Contig1” to “chrMT_Contig1,+,chrMT” to rename the sequence name for the mitochondrial genome assembly.

iii) Once all the modifications have been specified, run the following script to generate the final genome assembly as well as the associated genome-wide dotplot (Fig. 3b) and assembly statistics (Table 1):

$ bash LRSDAY.06.Supervised_Final_Assembly.2.sh

34) Re-run centromere annotation for the final genome assembly:

$ cd./../07.Centromere_Annotation

$ bash LRSDAY.07.Centromere_Annotation.sh

35) Annotate protein-coding genes and tRNAs for the final genome assembly. This step can be run with multiple threads.

$ cd./../08.Gene_Annotation

$ bash LRSDAY.08.Gene_Annotation.sh

**▴CRITICAL STEP** When running LRSDAY with your own data, if you have native transcriptome/EST assembly for your sequenced strain/species, you can first make a copy of the default configuration file “$LRSDAY_HOME/misc/maker_opts.ctl” and place it in the task-specific directory “$LRSDAY_HOME/Project_abc/08.Gene_Annotation”. Then, edit the line 16 of the configuration file that you have just created to provide the full path of your transcriptome/EST assembly. Finally, edit the line 13 of the bash script “LRSDAY.08.Gene_Annotation.sh” to provide the full path of your edited configuration file. This path should be: “$LRSDAY_HOME/Project_abc/08.Gene_Annotation/maker_opts.ctl”. For all these changes, you should replace “Project_abc” with the real directory name that you created in Step 24.

36) (Optional) A file containing a list of genes with suspicious annotations will be generated in the file “SK1.maker.manual_check.list”. As labeled in this file, these annotated gene models could be fragmented, frameshifted or containing internal stop codons. Potentially, these genes could be good candidates for pseudogenes. You can manually inspect the annotated gene models of these genes by loading the annotation result “SK1.maker.gff3” together with the protein/EST-alignment evidence files generated during the annotation (SK1.protein_evidence.gff3 and SK1.est_evidence.gff3) into IGV^22^ for further curation.

37) (Optional) Dedicated mitochondrial protein-coding and RNA annotation. If you are interested in studying mitochondrial genomes, we highly recommend running dedicated mitochondrial feature annotation with specialized software such as MFannot (http://megasun.bch.umontreal.ca/cgi-bin/mfannot/mfannotInterface.pl). Although MFannot does not support local installation by far, you can run MFannot conveniently via its web portal. Select the correct genetic code table (e.g. “3 Yeast Mitochondrial” for annotating yeast mitochondrial genomes) for your analysis.

38) Annotate transposable elements (TEs) for the final genome assembly. This step can be run with multiple threads.

$ cd./../09.TE_Annotation

$ bash LRSDAY.09.TE_Annotation.sh

39) (Optional) TE activity can be highly dynamic in the genome with many complex cases such as fragmentation and nested insertion. In LRSDAY, we used REannotate^23^ to automatically resolve these complex cases, which works greatly for most cases but it could occasionally misjoin two adjacent TEs when they are closely spaced. You can further inspect and curate the LRSDAY TE annotation (“SK1.TE.gff3”) by visualizing it in IGV^22^ together with the raw REannotate annotation (“SK1.REannotate.gff”). For each TE found in the file “SK1.TE.gff3”, examine its corresponding LTR and internal region structure based on the file “SK1.REannotate.gff” to check for misjoinings. If needed, you can manually edit the corresponding annotation in the “SK1.TE.gff3” file to decouple the misjoinings.

40) Annotate yeast telomere-associated core X elements for the final genome assembly:

$ cd./../10.Core_X_Element_Annotation

$ bash LRSDAY.10.Core_X_Element_Annotation.sh

41) (Optional) In LRSDAY, we label the identified core X elements to be “partial” if they are shorter than 300 bp. This should work for most cases but we recommend to inspect the generated alignment file (“SK1.X_element.aln.fa”) for further curation and to manually adjust such “partial” labeling in the annotation file (“SK1.X_element.gff3”) when needed.

42) Annotate yeast telomere-associated Y’ elements for the final genome assembly:

$ cd./../11.Y_Prime_Element_Annotation

$ bash LRSDAY.11.Y_Prime_Element_Annotation.sh

43) (Optional) Like for the core X element annotation, we used a hard length cutoff (3500 bp) to label if the identified Y’ elements are “partial”. We suggest to manually inspect the generated alignment file (“SK1.Y_prime_element.aln.fa”) to check if the “partial” labeling is needed and edit the annotation file (“SK1.Y_prime_element.gff3”) accordingly.

44) Orthology identification for protein coding genes. In this step, we create a gene orthology relationship list between the annotated proteome and the SGD *S. cerevisiae* reference proteome based on both sequence similarity and synteny conservation. Based on this list, we further attach SGD systematic names to our gene annotation as shown in the “Name=” field of the generated GFF3 file (“SK1.updated.gff3”). For a given annotated gene, when more than one orthologous gene can be found in the SGD reference proteome, we will labeled all of its co-orthologs in the “Name=” filed with “/” in between, whereas when no orthologous gene can be found, we will label its gene name as “Name=NA”. This step can be run with multiple threads.

$ cd./../12.Gene_Orthology_Identification

$ bash LRSDAY12.Gene_Orthology_Identification.sh

45) Integrate the annotation of different genomic features into a unified GFF3 file.$ cd./../13.Annotation_Integration $ bash LRSDAY.13.Annotation_Integration.sh

**? TROUBLESHOOTING**

Troubleshooting advice can be found in Table 2.

**TABLE 2.**
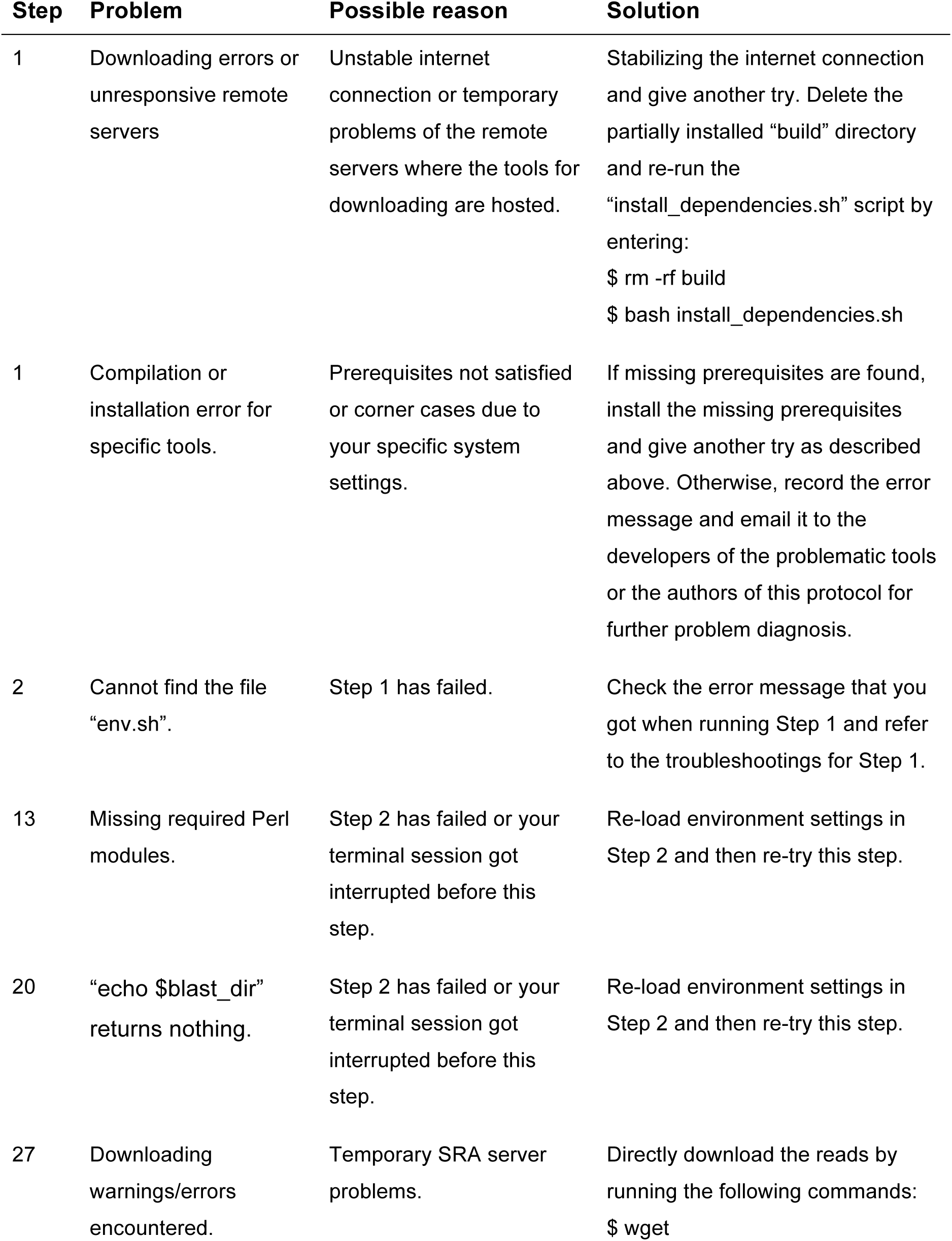
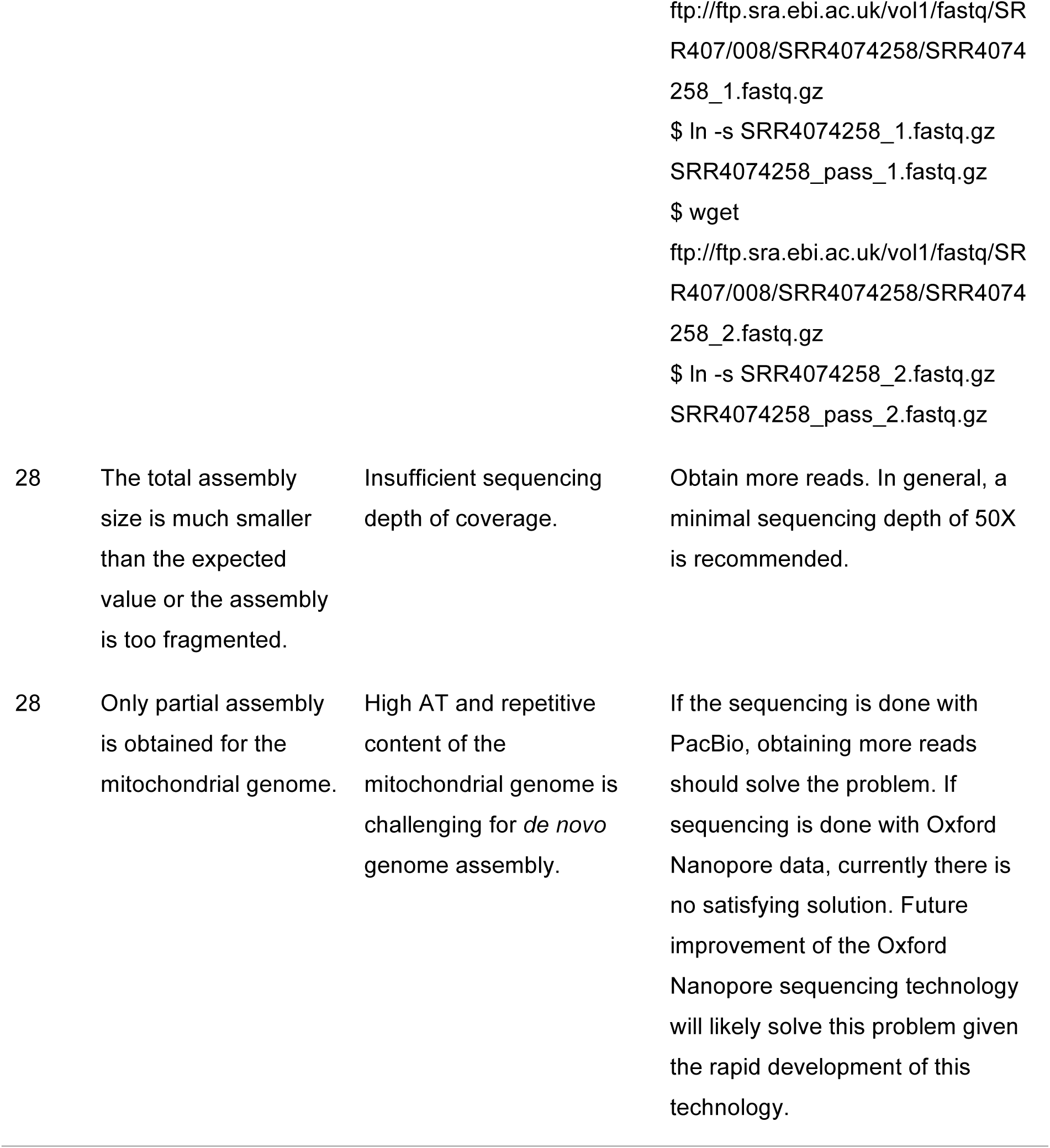
| Troubleshooting table.

## •**TIMING**

The following timing information was measured on a Linux computing server with an Intel Xeon CPU E5-2630L v3 (1.80GHz) using four threads. Enable multithreading can substantially decrease the processing time. Those optional steps were not processed when measuring the computation time.

Step 1-22, set up LRSDAY: < 1h.

Step 23-24, prepare the project directory for the test example: 5 s.

Step 25, prepare the reference genomes for the test example: 1 s.

Step 26, prepare the long reads for the test example: 22 min.

Step 27, prepare the Illumina reads for the test example: 20 min.

Step 28, *de novo* assembly using long-reads: 18 h.

Step 29, assembly polishing using Illumina reads: 1h.

Step 30, reference-guided scaffolding for the raw assembly: 8 min.

Step 31, centromere identity profiling for the scaffolded assembly: 1 s.

Step 32, mitochondrial genome assembly improvement: 4 min.

Step 33, finalize genome assembly with supervised editing: 1 min.

Step 34, centromere annotation for the final assembly: 1s.

Step 35, protein-coding gene and tRNA annotation for the final assembly: 21 h.

Step 38, TE annotation for the final assembly: 5 min.

Step 40, core X element annotation for the final assembly: 2 min.

Step 42, Y’ element annotation for the final assembly: 2.5 min.

Step 44, gene orthology identification for the annotated protein-coding genes: 20 min.

Step 45, final annotation integration: 1 min.

### ANTICIPATED RESULTS

In this section, we describe the major outputs of the test run “Project_Example” as follows. The final genome assembly and annotation outputs generated with this test example is provided in the directory “Example_Outputs” for users to make direct comparison with their own results.

Task 01 (Step 28):

- SK1.canu.fa: the long-read-based *de novo* genome assembly containing all the contigs assembled by Canu^12^.
- SK1.canu.stats.txt: the summary table of basic assembly statistics, including information on the number of the assembled sequences, the total length of the assembled sequences, the minimal, maximal, mean and median lengths of the assembled sequences, the N50, L50, N90, and L90 of the assembled sequences, as well as the base composition (A%, T%, G%, C%, AT%, GC% and N%) of the assembled sequences. Users can compare this file with the similar file generated in Step 30 and Step 33. We also summarized such comparison in Table 1.
- SK1_canu_out: the directory containing all the output files of Canu.

Task02 (Step 29):

- SK1.pilon.fa: the polished genome assembly generated by Pilon^24^.
- SK1.pilon.vcf: the variants identified by Pilon based on short reads mapping against the input genome assembly.
- SK1.pilon.changes: a space-delimited record of all the changes that Pilon made during the assembly polishing. The four columns are: the original sequence coordinate, the new sequence coordinate after the correction, the original base, the new base after the correction.
- SK1.realn.bam: the bam file of short reads mapping against the input genome assembly.

Task 03 (Step 30):

- SK1.ragout.fa: the scaffolded genome assembly based on the reference genome.
- SK1.ragout.stats.txt: the summary table of basic assembly statistics for the scaffolded genome assembly. Users can compare this file with the similar file generated in Step 28 and Step 33. We also summarized such comparison in Table 1.
- SK1.ragout.agp: the NCBI AGP file recording the order and orientation of each input contigs used during scaffolding.
- SK1_ragout_out: the directory containing all the output files of Ragout^25^.
- SK1.ragout.filter.pdf: the genome-wide dotplot for the comparison between the scaffolded assembly and the reference genome.
- SK1.mt_contig.list: the list of contigs in the scaffolded assembly that corresponds to the mitochondrial genome. This file will be used for Step 32.
- SK1.mt_contig.fa: the sequences of the contigs in the scaffolded assembly that corresponds to the mitochondrial genome.
- SK1.ragout.chrMT.filter.pdf: the dotplot for the comparison between the scaffolded itochondrial genome assembly and the reference mitochondrial genome.

Task 04 (Step 31):

- SK1.centromere.gff3: the profiled centromere idenities for the scaffolded genome assembly.

Task 05 (Step 32):

- SK1.mt_improved.fa: the improved genome assembly with better processing (re-assembling and circularization) of the mitochondrial genome.
- SK1.mt_improved.chrMT.filter.pdf: the dotplot for the comparison between the improved mitochondrial genome assembly and the reference mitochondrial genome. You should see much better collinearity in this plot than the similar plot generated in Step 30.

Task 06 (Step 33):

- SK1.modification.list: the assembly modification list for manual editing to guide the final genome assembly.
- SK1.final.fa: the final genome assembly generated by LRSDAY.
- SK1.final.filter.pdf: the genome-wide dotplot for the comparison between the final genome assembly and the reference genome.
- SK1.final.stats.txt: the summary table of basic assembly statistics for the final genome assembly. Users can compare this file with the similar file generated in Step 28 and Step 30. We also summarized such comparison in Table 1.

Task 07 (Step 34):

- SK1.centromere.gff3: the centromere annotation for the final genome assembly.

Task 08 (Step 35):

- SK1.maker.raw.gff3: the raw MAKER^21^ annotation for protein-coding genes and tRNA genes.
- SK1.maker.gff3: the processed MAKER annotation for protein-coding genes and tRNA genes with systematically assigned gene IDs.
- SK1.maker.output: the output directory of the MAKER run.
- SK1.protein_evidence.gff3: the protein-to-genome alignment evidences generated by MAKER. This file can be used for manual curation of those suspicious annotations.
- SK1.est_evidence.gff3: the EST-to-genome alignment evidences generated by MAKER. This file can be used for manual curation of those suspicious annotations.
- K1.maker.trimmed_cds.fa: the CDSs of the MAKER protein-coding annotation with the out-of-frame parts trimmed.
- SK1.maker.trimmed_cds.log: the log file of the CDS trimming for the MAKER protein-coding gene annotation.
- SK1.maker.pep.fa: the translated protein sequences of the trimmed CDSs derived from the MAKER protein-coding gene annotation.
- SK1.maker.manual_check.list: a list of suspicious gene annotations for manual curation.
- SK1.maker.PoFF.gff: the gene synteny information derived from “SK1.maker.gff3”, which will be used for Task 12 (Step 44).
- SK1.maker.PoFF.ffn: same as “SK1.maker.trimmed_cds.fa” but with simpler sequence IDs, which could be used for Task 12 (Step 44).
- SK1.maker.PoFF.ffa: same as “SK1.maker.protein.fa” but with simpler sequence IDs, which will be used for Task 12 (Step 44).

Task 09 (Step 38):

- SK1.REannotate.gff: the raw TE annotation from REannotate. This file can be used for further curating TE annotation.
- SK1.TE.gff3: the final TE annotation from LRSDAY.

Task 10 (Step 40):

- SK1.X_element.gff3: the final core-X element annotation from LRSDAY.
- SK1.X_element.fa: the sequences of all the annotated core X elements.
- SK1.X_element.aln.fa: the sequence alignment of all the annotated core X elements for further checking whether the annotated feature is complete or partial and whether this is consistent with the labeling in the annotation file “SK1.X_element.gff3”.

Task 11 (Step 42):

- SK1.Y_prime_element.gff3: the final Y’ element annotation from LRSDAY.
- SK1.Y_prime_element.fa: the sequences of all the annotated Y’ elements.
- SK1.Y_prime_element.aln.fa: the sequence alignment of all the annotated Y’ elements for further checking whether the annotated feature is complete or partial and whether this is consistent with the labeling in the annotation file “SK1.Y_prime_element.gff3”.

Task 12 (Step 44):

- SK1.proteinortho: the gene orthology mapping between the annotated assembly and the reference genome based only on sequence similarity.
- SK1.poff: the gene orthology mapping between the annotated assembly and the reference genome based on both sequence similarity and synteny conservation.
- SK1.updated.gff3: updated gene annotation with reference-based gene name labeling.

Task 13 (Step 45):

- SK1.final.gff3: the final integrated annotation from LRSDAY.
- SK1.final.trimmed_cds.fa: the CDSs of the final protein-coding gene annotation with the out-of-frame parts trimmed.
- SK1.final.trimmed_cds.log: the log file of the CDS trimming for the final protein-coding gene annotation.
- SK1.final.pep.fa: the translated protein sequences of the trimmed CDSs derived from the final protein-coding gene annotation.
- SK1.final.manual_check.list: a list of suspicious gene annotations for manual curation.

## AKNOWLEDGEMENTS

We thank Gilles Fischer, Samuel O’Donnell and Lorenzo Tattini for testing LRSDAY and providing valuable feedback. This work was supported by ATIP-Avenir (CNRS/INSERM), Fondation ARC pour la Recherche sur le Cancer (PJA20151203273), Marie Curie Career Integration Grants (322035), Agence Nationale de la Recherche (ANR-16-CE12-0019, ANR-13-BSV6-0006-01 and ANR-11-LABX-0028-01), Cancéropôle PACA (AAP émergence 2015) and a DuPont Young Professor Award to G.L. J.-X.Y. is supported by a postdoctoral fellowship from Fondation ARC pour la Recherche sur le Cancer (PDF20150602803).

## AUTHOR CONTRIBUTIONS

J.-X.Y. designed, implemented, and tested the LRSDAY workflow. G.L. coordinate the work. J.-X.Y. and G.L. wrote the manuscript.

## BOXES

#### BOX1: Pre-shipped supporting data for LRSDAY

With LRSDAY, we pre-shipped the following supporting datasets for the automatic execution of LRSDAY. Unless labeled otherwise, most of these pre-shipped datasets were either described or generated in our previous study^5^.

ATP6.cds.fa # The coding sequence (CDS) of the *S. cerevisiae* S288C *ATP6* gene. evm_weights.txt # the evidence weights file for EVidenceModeler (EVM)^26^ (shipped with MAKER).

FungiDB_Sc.est.simple.fa # the *S. cerevisiae* EST data from FungiDB (http://fungidb.org/fungidb/).

fuzzy_defragmentation.txt # the fuzzy defragmentation file for REannotate^23^. Proteome_DB_for_annotation.CDhit_I95.fa # our curated proteome dataset for *S. cerevisiae* and other closely related yeast *sensu stricto* species.

query.Y_prime_element.long.fa # the sequence of a representative *S. cerevisiae* Y’ element. S288C.ASM205763v1.fa.gz # the *S. cerevisiae* S288C genome assembly.

S288C.ASM205763v1.noncore_masked.fa.gz # the *S. cerevisiae* S288C genome assembly with subtelomeres and chromosome-ends hard-masked.

S288C.centromere.fa # the centromere sequence of *S. cerevisiae* S288C.

S288C.gene.hmm # the hidden Markov model (HMM) for *de novo* gene annotation based on *S. cerevisiae* S288C.

S288C.X_element.hmm # the hidden Markov model (HMM) for the core X element annotation based on *S. cerevisiae* S288C.

Sc-meth.sites # the *S. cerevisiae* methylation sites (shipped with snoScan^27^). Sc-rRNA.fa # the *S. cerevisiae* rRNA sequences (shipped with snoScan^27^).

SGDref.PoFF.faa # the proteinortho proteome file generated for the SGD reference genome. SGDref.PoFF.ffn # the proteinortho CDS file generated for the SGD reference genome.

SGDref.PoFF.gff # the proteinortho gene order gff file generated for the SGD reference genome.

te_proteins.fasta # protein sequences for genes encoded within TEs (shipped with MAKER^21^). TY2_specific_region.fa # the sequence of a Ty2 specific regions for differentiating Ty1 and Ty2. TY_lib.Yue_et_al_2017_NG.fa # a custom RepeatMasker library for Ty annotation in *S. cerevisiae* and *S. paradoxus*.

TY_lib.Yue_et_al_2017_NG.LTRonly.fa # representative Ty LTR sequences of *S. cerevisiae* and *S. paradoxus*.

#### BOX2: Tips for adapting LRSDAY to other eukaryotic organisms

The backbone modules of LRSDAY can be easily adapted for other eukaryotic organisms. Here are some tips with regard to this:

1) For Task 01, be sure to adjust the genome size parameter in line 13 of the bash script “LRSDAY.01.Long-read-based_Genome_Assembly.sh” based on the estimated genome size of the organism that you sequenced.

2) For Task 03, be sure to modify the bash script “LRSDAY.03.Reference-guided_Assembly_Scaffolding.sh” to provide the reference genome file of your sequenced organisms to guide the scaffolding and chromosome assignment. It is very likely that the chromosomal cores and subtelomeres of your reference genome have not been clearly defined. In such case, you can provide the same reference genome file for both the “ref_genome_raw=” and “ref_genome_noncore_masked=” parameters. The “chrMT_tag=” and “gap_size=” parameters should also be adjusted based on your own project.

3) By default, Task 03 performs reference-guided scaffolding using the whole genome alignment constructed by Sibelia^28^. Sibelia is not designed for processing small genomes (<100 Mb). For organisms with large genomes, you should install Progressive Cactus (https://github.com/glennhickey/progressiveCactus) and use it to build the whole genome alignment in HAL format and feed it directly to Ragout^25^. Please refer to Ragout’s user manual for this advanced usage. We have preshipped the HAL tools (https://github.com/ComparativeGenomicsToolkit/hal) to enable Ragout to process the HAL file generated by Progressive Cactus.

4) For Task 05, be sure to modify the “gene_start=”, “ref_genome_raw=”, and “chrMT_tag=”parameters in the bash script LRSDAY.05.Mitochondrial_Genome_Assembly_Improvement.sh” based on your own project.

5) While many of the genomic feature annotation tasks are yeast-specific, Task 08 (protein-coding genes and tRNA annotation) can be adapted for any eukaryotic organism. Please refer to MAKER’s own Wiki page (http://weatherby.genetics.utah.edu/MAKER/wiki/index.php/Main_Page) and protocols (https://www.ncbi.nlm.nih.gov/pmc/articles/PMC4286374/) for technical details and advanced usage.

6) While Task 09 (TE annotation) is heavily tuned for yeasts, the same tools that we used here (RepeatMasker^20^ and REannotate^23^) can be used for any eukaryotic organism. We recommend to read their respective manuals for adapting these tools in your own study.

7) For Task 12, you need to edit the “ref_PoFF_faa=” and “ref_PoFF_gff=” parameters based on the reference gene annotation that you used. Please check ProteinOrtho’s manual (https://www.bioinf.uni-leipzig.de/Software/proteinortho/manual.html) for more details on required file formats. The pre-shipped Perl script “prepare_PoFFfaa_simple.pl” and “prepare_PoFFgff_simple.pl” in the “$LRSDAY_HOME/scripts” directory should help for this.

